# Estrogen Receptor Beta Activation Coordinates Liver Lipid Remodeling and Metabolic Fluxes, Preventing Lipotoxicity

**DOI:** 10.1101/2025.08.14.670313

**Authors:** Debora Santos Rocha, Eloisa Vilas-Boas, Camille C. Caldeira da Silva, Everton L. Vogt, Marcos Yoshinaga, Mariana Pacheco Teixeira de Carvalho, Maiara I. C. Queiroz, Tiago Eugênio Oliveira da Silva, Sayuri Miyamoto, Alicia J. Kowaltowski

## Abstract

Non-selective hormone replacement with estradiol is known to improve metabolic homeostasis during menopause. However, this treatment is not recommended for individuals with genetic predisposition to hormone-responsive cancers. In contrast, selective activation of estrogen receptor beta (ERβ) has shown promising results, promoting antitumor effects and modulating metabolic outcomes, although the mechanisms in which these changes occur remain poorly understood. We investigated the effects of ERβ activation using diarylpropionitrile (DPN), a selective ERβ agonist, in both an *in vivo* model of post menopause and *in vitro* model of metabolic overload. Female Wistar rats were submitted to ovariectomy (OVX) and later treated with DPN. ERβ agonist treatment recovered fasting glucose and lipid profiles, improved pancreatic islet morphology, and reduced retroperitoneal white adipose tissue. Serum ketone bodies and free fatty acid levels were also recovered to control levels, suggesting modulation in liver lipid oxidation. DPN treatment also promoted substantial lipidome remodeling specifically in OVX animals, generating a lipid-buffering profile. To isolate the effects mechanistically, hepatocytes were submitted to nutrient overload and treated with DPN. *In vitro*, DPN also recovered ketone body secretion and promoted an increased dependence on complete fatty acid oxidation, as well as decreased metabolic flexibility, as assessed by modulated extracellular flux analysis. Overall, these findings demonstrate a new role of ERβ in the modulation of hepatic lipid and ketone metabolism, with positive metabolic outcomes in estradiol-deficient animals.

## 1. INTRODUCTION

Estrogen signaling plays a central role not only in reproductive functions, but also in energy metabolism, stimulating insulin production and secretion in pancreatic beta cells, and improving peripheral insulin signaling in the adipose tissue, muscle, and liver (1,2). Accordingly, postmenopausal decreases in estradiol levels lead to various metabolic consequences, including increased risk of metabolic disease development. This is observed both in humans and in animal models of estrogen deficiency induced by ovariectomy or hypothalamic-pituitary-gonad axis modulation (3). Estrogen deficiency also promotes the accumulation of visceral white adipose tissue and circulating free fatty acid (FFA) overload (4,5). Chronically increased FFA levels compromise postprandial muscle glucose uptake and promote decompensation of hepatic glucose production. Consequent long-term glucotoxicity and lipotoxicity affect endocrine pancreatic responses to energy demands, culminating in defective insulin production and secretion, worsening the already impaired peripheral tissue insulin sensitivity (6,7).

In this sense, it is reasonable to assume that hormone replacement therapy (HRT) with estradiol would promote metabolic benefits postmenopause. In fact, HRT modulates adipose tissue redistribution to an insulin-sensitive phenotype, increasing subcutaneous and decreasing visceral depots (8), in addition to attenuating the metabolic imbalance associated with type II diabetes mellitus (9,10). However, non-selective activation of estradiol receptors (ERs) often provides only transient benefits which wane over time, and displays risks such as increased incidence of hormone-responsive tumor development in patients with gynecological cancer predisposition (11,12).

Estradiol used in HRT activates the 3 main types of estrogen receptors: nuclear receptors alpha (ERα) and beta (ERβ) as well as membrane receptors (GPER). ERα and ERβ trigger estrogen response element binding, modifying gene transcription and, consequently, protein synthesis. These genomic responses to estrogens occur within hours. Conversely, the activation of GPER leads to a rapid response, within seconds or minutes, modulating cytosolic signaling processes including calcium influx (13) and activation of PKA and PKC pathways (14).

Selective ERα activation attenuates energy imbalance, improves insulin sensitivity, changes white adipose tissue (WAT) phenotypes and promotes browning (2,15,16). The outcomes of selective ERβ activation results are still poorly characterized, although data also suggest changes in WAT storage patterns, shifting from visceral to subcutaneous predominance (15), as well as WAT browning, and increased insulin sensitivity signs (17) as evidenced by enhanced glucose uptake in muscle fibers (18). These effects remain to be characterized mechanistically.

In the liver, estradiol reduces the imbalanced accumulation of triacylglycerol, and consequently steatosis, while also reducing gluconeogenesis, results that are generally attributed to ERα activation due to its relative high expression levels (19–21). The selective activation of ERβ in female rodents is also capable of triggering responses in hepatocytes, reducing adipogenesis and lipid accumulation (22). If these effects are also observed under postmenopausal conditions remains to be explored.

Determining if selective ERβ activation promotes metabolic changes postmenopause is of interest because, while ERα activation is associated with the development of hormone-dependent cancers (23), ERβ activation has antitumor (24–26), cell cycle inhibitory (27), and antiangiogenic effects (23,28). It is thus reasonable to assume that selective modulation of ERβ would be a promising alternative therapeutic strategy for postmenopausal metabolic syndrome. Therefore, the aim of this paper is to describe the metabolic effects of selective beta estradiol receptor (ERβ) activation under postmenopausal conditions, with a focus on the liver as a central metabolic hub.

## 2. MATERIALS AND METHODS

### 2.1. Animals

Female Wistar rats (n = 5-7 per group, total = 26) were randomized, acclimatized for 20 days, and maintained at controlled temperatures (23°C) with 12-12 h light/dark cycles and *ad libitum* access to food/water. A pilot and an experimental study (including all experimental groups) were conducted. All procedures were conducted in accordance with the Ethical Principles of Animal Manipulation of the local animal ethics committee (CEUA-IQ/USP protocol 244/2022), the National Research Council Guide for the Care and Use of Laboratory Animals, and follow ARRIVE Guidelines (29–31).

At 90 days of age, animals were submitted to ovariectomy as described in the “Ovariectomy surgery” section. Fifteen days later, they began treatment with ERβ agonist diarylpropionitrile (DPN, 1 mg. kg^-1^ .day^-1^, s.c.) or vehicle (sesame oil) for 15 days. Four groups were studied: sham surgery treated with vehicle (Vehicle); sham surgery treated with DPN (DPN); ovariectomized animals treated with vehicle (OVX vehicle); and ovariectomized animals treated with DPN (OVX DPN). The animals were observed for 30 days for their food intake, weight gain, and glucose tolerance (described under “Glucose tolerance test”). At the endpoint, animals were anesthetized and killed by cardiac punction. Serum was obtained after total blood centrifugation (20 minutes at 300 g) and stored for further analysis. Retroperitoneal white adipose tissue (WATr), uteruses, and livers were collected and weighed. WATr, pancreases, and livers were evaluated for histological characteristics (see “Histology analysis”). The liver was also used for mitochondrial isolation; fresh mitochondria were used to measure oxygen consumption rates as described in the “Oxygen consumption assay” section.

### 2.2. Ovariectomy surgeries

90-day-old female Wistar rats were anesthetized (4% isoflurane for induction and 1.5% for maintenance) and monitored for temperature, heartbeats, and ventilation rates. Rats were positioned in ventral decubitus, and an incision was made between the lower costal margin and thigh, approximately 1 cm from the midline. The muscle wall was retracted, then the ovary was located within the adipose tissue and removed after ligation of the corresponding uterine tube by sectioning between the ligature and the ovary. Muscle and skin were then sutured, and the procedure was repeated on the contralateral side (32). After surgery, the animals were monitored and treated with tramadol (20 mg/kg) for analgesia. None of the animals presented pain or distress considering the Grimace scale (33) Sham surgery animals also underwent the same procedures, except for the removal of the ovaries.

### 2.3. Glucose tolerance tests

Glucose tolerance was evaluated before the ovariectomy surgery (week 1); 15 days after surgery and before DPN treatment (week 3); and after 30 days of surgery and 15 days of DPN treatment (week 5). The animals were submitted to 6 hours of food deprivation and basal glucose levels were registered. Glucose (1 g per kg of body weight) was diluted in isotonic saline solution (NaCl 0.9%) and injected intraperitoneally. Blood glucose levels were registered at 5, 15, 30, 60, 90, and 120 minutes using a Roche Accu-Chek glucometer.

### 2.4. Histological analysis

WATr, pancreas, and liver tissues were fixed in buffered formalin, dehydrated with ethanol solutions (70 to 100 %), embedded in paraffin, sectioned, mounted on slides, deparaffinized, and stained with hematoxylin/eosin. Images were registered with an Axio Scan.Z1 slide scanner and analyzed using QuPath software.

Pancreases were evaluated for islet morphometric parameters. The endocrine tissue was visually identified, and islets were circulated individually. The shape descriptor tool was used to measure size and circularity. Islets were divided into quartiles as small, medium, or large using the control group as a reference (34–36).

WATr was analyzed according to Palomäki *et al.* (37). The software was trained using the “train pixel classifier” tool. The brush and expected size range were used as training inputs. The “create objects” tool was then used and cell-by-cell visual confirmation ensured correct output compared to the original image. The “shape descriptors” tool was used to analyze average adipocyte areas (µm^2^).

Liver histology was evaluated for macrovesicular steatosis, considering the non-stained area within the tissue (apart from the portal vein branch area). Positive areas for macrovesicular steatosis were described as percentage of total area and interpreted as neutral lipid accumulation *in vivo* (38,39).

### 2.5. Biochemical serum analysis

Serum total cholesterol, HDL, LDL, triglycerides, free fatty acids, and beta hydroxybutyrate were detected using commercial kits (LabTest^®^, FUJIFILM^®^, and Sigma Aldrich^®^). VLDL levels were calculated by subtracting the measured HDL and LDL cholesterol fractions from total cholesterol. Sample preparation, dilution, and standard curve protocols were conducted according to manufacturer instructions.

### 2.6. Tissue glycogen and triglycerides

Samples were hydrolyzed with 30% KOH at 100°C. Glycogen was precipitated in 70% ethanol, washed with distilled water, and converted into glucose by adding 4 M HCl at 100°C. Glycogen levels were measured based on glucose quantities resulting from hydrolysis compared to a commercial standard glycogen sample (Sigma-Aldrich) previously hydrolyzed in HCl. Detection was performed using a commercial glucose assay kit (LabTest^®^). Triacylglycerol was detected using a commercial kit (LabTest^®^). Data were expressed as µg triglycerides per µg protein and mg glycogen per mg tissue.

### 2.7. Lipidomics

Liver samples (approximately 200 mg) were homogenized in PBS buffer containing 20 µM deferoxamine and subjected to methyl-tert-butyl ether (MTBE) extraction (40). Lipid extracts were analyzed via an untargeted lipidomics approach using ultra-high-performance liquid chromatography (UHPLC; Nexera, Shimadzu, Kyoto, Japan) coupled with electrospray ionization tandem time-of-flight mass spectrometry (ESI-MS/MS) (Triple TOF® 6600, Sciex, Concord, USA). Lipid species were identified by inspecting MS/MS spectra using PeakView® (Sciex). Each lipid molecular species was quantified by normalizing the precursor ion peak area to that of the corresponding internal standard using MultiQuant® (Sciex). Concentrations were calculated based on the ratio of integrated MS data to the sample volume, and results were expressed as a percentage of total lipids or in nmols per mg of protein. The coefficient of variation (CV) was calculated to filter lipid species displaying CV>20%, based on 5 quality control samples measured during the experimental routines in ESI-MS/MS. Data was log10-transformed to achieve normal distribution. One-way ANOVA followed by FDR correction (196 altered compounds, p<0.05) was used to compare the amount of lipids changed between four experimental groups. A heatmap displaying the top 75 lipid species (lowest ANOVA p-values) were built using Euclidean distance and Ward clustering. The top 75 lipid species (lowest ANOVA p-values) are displayed. For Principal Component Analysis (PCA) analysis, normalized (log10-transformed) data were visualized in a 2D PCA plot showing PC1 and PC2 (X% and Y% of variance, respectively). For cholesteryl esters (CE) and triglycerides (TG), only CE and TG associated lipids with statistically significant differences (t-test, two-group comparison; p<0.05) were analyzed; fold-change was then calculated for each significantly altered feature. All statistical analysis were performed in MetaboAnalyst (https://www.metaboanalyst.ca/,(41)). For TG fatty acid (FA) analysis, the molar abundance of each individual fatty acid (FA) within TG species was calculated and normalized to the total TG molar concentration for t-test and fold-change analyses. TG were further categorized by FA saturation status as: saturated (S), monounsaturated (M), polyunsaturated (P, 2-3 double bonds), and highly unsaturated (H, >3 double bonds).

### 2.8. Mitochondrial oxygen consumption measurements

The liver was chopped, suspended in isolation buffer (250 mM sucrose, 10 mM HEPES, 1 mM EGTA, 1 mM EDTA, 1 mg/mL BSA, pH 7.2, at 4°C), and manually homogenized using a Potter–Elvehjem tissue grinder. The suspension was centrifuged twice (800 g, 4°C, 4 min), and the pellet discharged. The supernatant was further centrifuged at 9000 g, 4°C for 10 min. The pellet was resuspended in resuspension buffer (300 mM sucrose, 10 mM HEPES, and 2 mM EGTA, pH 7.2) and centrifuged again (12000 g, 4°C, 10 min). The pellet was resuspended in 1 mL resuspension buffer, and total protein was quantified using the Bradford method.

For respirometry assays, 250 µg or 50 µg protein (for CI or CII–linked respiration, respectively) of isolated mitochondria were incubated in a buffer containing 120 mM sucrose, 65 mM KCl, 2 mM MgCl₂, 1 mM KH₂PO₄, 1 mM EGTA, 10 mM HEPES, and 0.1% fatty acid-free BSA, pH 7.2, at 37°C in an Oroboros O2k high-resolution respirometer (Innsbruck, Austria). 1 mM pyruvate plus 1 mM malate and 1 mM α-KG, or 1 mM succinate plus 1 μM rotenone were used to induce CI-linked and CII-linked oxygen consumption, respectively. Oxygen consumption was monitored through sequential additions of ADP (1 mM) to induce state 3, oligomycin (1 µM) to induce state 4, and CCCP (titrated in 100 nM additions) for state 3u, as described in previous studies (42).

### 2.9. Cell cultures

Non-tumor alpha mouse liver 12 cells (AML12, ATCC® CRL-2254™) were cultivated in Dubelco’s modified Eagle’s minimal essential medium (DMEM, GIBCO^®^, Life Technologies) - low glucose (5.6 mM) - supplemented with 10% fetal bovine serum (FBS) and 1% antibiotic-antifungal solution (Microlab^®^), 1.7 µM Insulin, 0.069 µM Transferrin, 0.038 Sodium Selenite, without Phenol Red. The cells were kept in a humidified incubator at 37°C, in an atmosphere of 5% CO_2_ and 95% air.

Cells were plated and, after 24 hours (∼80% confluence), the medium was replaced with treatment medium lacking FBS. Experimental groups were as follows: low glucose-BSA (LGB), high glucose-palmitate (HGP), and HGP plus ERβ agonist (DPN at 0.1 nM or 1 nM for an additional 24 h). The LGB group received BSA (0.25%) plus low glucose (5.6 mM). The HGP group was exposed to BSA-Palmitate (200 µM) plus high glucose (20 mM).

### 2.10. Fluorescence microscopies

24 h after treatment, mitochondrial morphology was verified in cell cultures through live 3D fluorescence microscopy. Cells were loaded with Mitotracker Deep Red (50 nM) and BODIPY (1 µM) for 20 minutes. The media was then replaced by PBS, and images of live cells were taken using a Leica DMi8 fluorescence microscope (43). The mitochondrial network was analyzed using the Mitochondrial Analyzer plugin (ImageJ software) and evaluated for mean volume and sphericity. Lipid droplets were assessed for their count and area per cell. 40 to 50 cells were quantified per group for each biological replicate. At least 5 different captured images were analyzed per biological replicate (n = 5).

### 2.11. Mitostress tests

After treatment, cells were washed in PBS twice and incubated in DMEM containing 5.6 mM glucose, 1% penicillin/streptomycin, and 5 mM HEPES, without FBS or bicarbonate. Oxygen consumption rates (OCRs) were quantified using an extracellular flux analyzer (Seahorse XF24). Basal respiration was registered, and mitochondrial modulators were injected sequentially to quantify ATP-linked, maximal, and non-mitochondrial respiration by injecting oligomycin (1 µM), CCCP (1 µM), and antimycin plus rotenone (1 µM each), respectively. All mitochondrial modulators were titrated separately in preliminary experiments under LGB and HGP conditions (due to the possible quenching effect of fatty acids), and doses used were the minimal required for a maximal response (42).

### 2.12. Metabolic flexibility assays

After treatment, cells were washed in PBS twice and incubated in DMEM containing 5.6 mM glucose, 1% penicillin/streptomycin, and 5 mM HEPES, without FBS or bicarbonate. OCRs were quantified using a Seahorse XF24 extracellular flux analyzer. In the metabolic flexibility assay, cells were sequentially submitted to specific oxidation pathway inhibitors: Etomoxir (CPT1/FFA oxidation inhibitor - 7 µM), BPTES (glutaminase/GSL1 inhibitor - 8 µM), and UK5099 (mitochondrial pyruvate transporter inhibitor - 6 µM). A decrease in OCR after the inhibitor injection indicates the dependence on the specific substrate (when only this pathway is inhibited), and the maximum capacity to oxidize the specific substrate is obtained when the alternative pathways are inhibited. Metabolic flexibility was calculated as the difference between dependence and capacity, indicating cell effectiveness in changing substrate use. All concentrations used were previously titrated under LGB and HGP conditions, as described above.

### 2.13. Western blots

Cells were homogenized in RIPA buffer containing 50 mM HEPES, 40 mM NaCl, 50 mM NaF, 2 mM EDTA, 10 mM sodium pyrophosphate, 10 mM sodium glycerophosphate, 2 mM sodium orthovanadate, 1% Triton-X 100, and protease inhibitors without EDTA, then centrifuged at 13,000 rpm for 10 minutes at 4°C. Lysates were diluted in Laemmli buffer, and proteins were separated using a 10% denaturing polyacrylamide gel. Proteins were transferred to nitrocellulose membranes and incubated with specific antibodies (**Table 1**). Secondary antibodies were added for visualization of bands, which were semi-quantified using densitometric analysis with Image J.

**Table 1.**
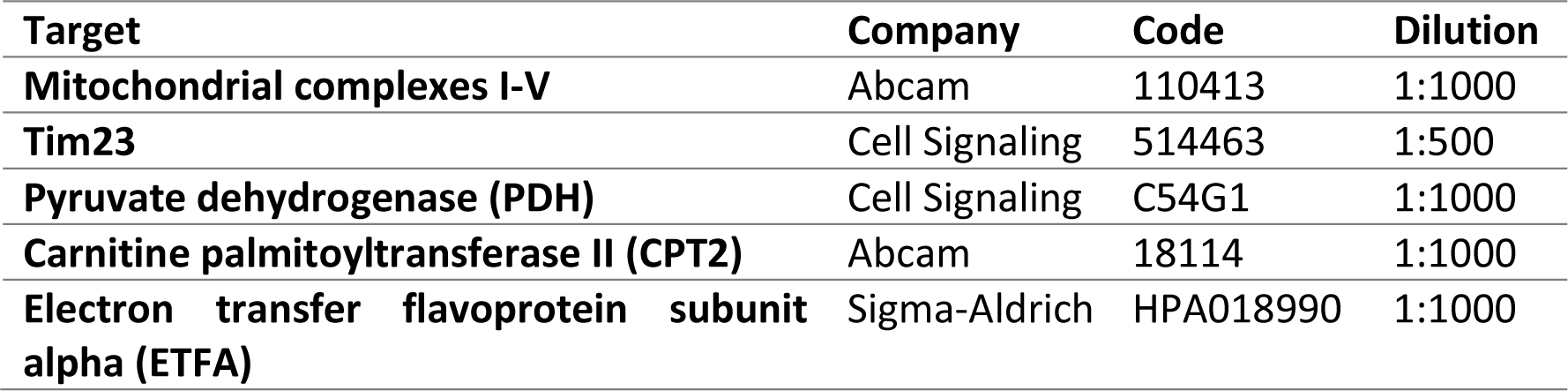
Antibodies and dilutions.

| **Target** | **Company** | **Code** | **Dilution** |
| --- | --- | --- | --- |
| **Mitochondrial complexes I-V** | Abcam | 110413 | 1:1000 |
| **Tim23** | Cell Signaling | 514463 | 1:500 |
| **Pyruvate dehydrogenase (PDH)** | Cell Signaling | C54G1 | 1:1000 |
| **Carnitine palmitoyltransferase II (CPT2)** | Abcam | 18114 | 1:1000 |
| **Electron transfer flavoprotein subunit alpha (ETFa)** | Sigma-Aldrich | HPA018990 | 1:1000 |

### 2.14. Statistical analysis

Data were evaluated in Prism Graphpad software, using ANOVA and Tukey posttests when data presented gaussian distribution and variance homogeneity, or correlate tests for non-gaussian distributions. For animal data, two-way (OVX and DPN variables) unpaired tests were used to compare different individuals, while repeated measures were applied to compare the same individual over time. The biological replicates necessary were based on the minimum sample size reported in prior studies assessing ovariectomy and subsequent biochemical analyses, ensuring statistical reliability of the results with minimal animal use (32,34,35,37). For *in vitro* data, one-way paired tests were conducted, since each biological replicate was divided across all experimental conditions. Also, unpaired data are displayed as individual dots, and paired analyses are plotted as continuous lines connecting individual experiments. All groups were verified statistically for outliers to be considered in the analysis. Significance was considered for differences where the p value was lower than 0.05.

## 3. RESULTS

### 3.1. ERβ agonist DPN modulates glucose homeostasis in ovariectomized (OVX) animals

The *in vivo* metabolic effects of OVX were monitored in female rats over the course of five weeks. Week 1 data were collected under baseline conditions, before the intervention protocol began, just prior to surgeries (OVX or sham). ERβ agonist (DPN) treatment began 15 days later, at Week 3. The experimental design is shown in **Figure 1A**. Measurements included body weight (BW), serum glucose levels, and glucose tolerance tests (GTT) repeated in Week 3 (15 days after OVX surgery, just prior to DPN treatment) and Week 5 (30 days after surgery and at 15 days of DPN treatment), when animals were euthanized and tissue analysis was conducted.

**Figure 1.**
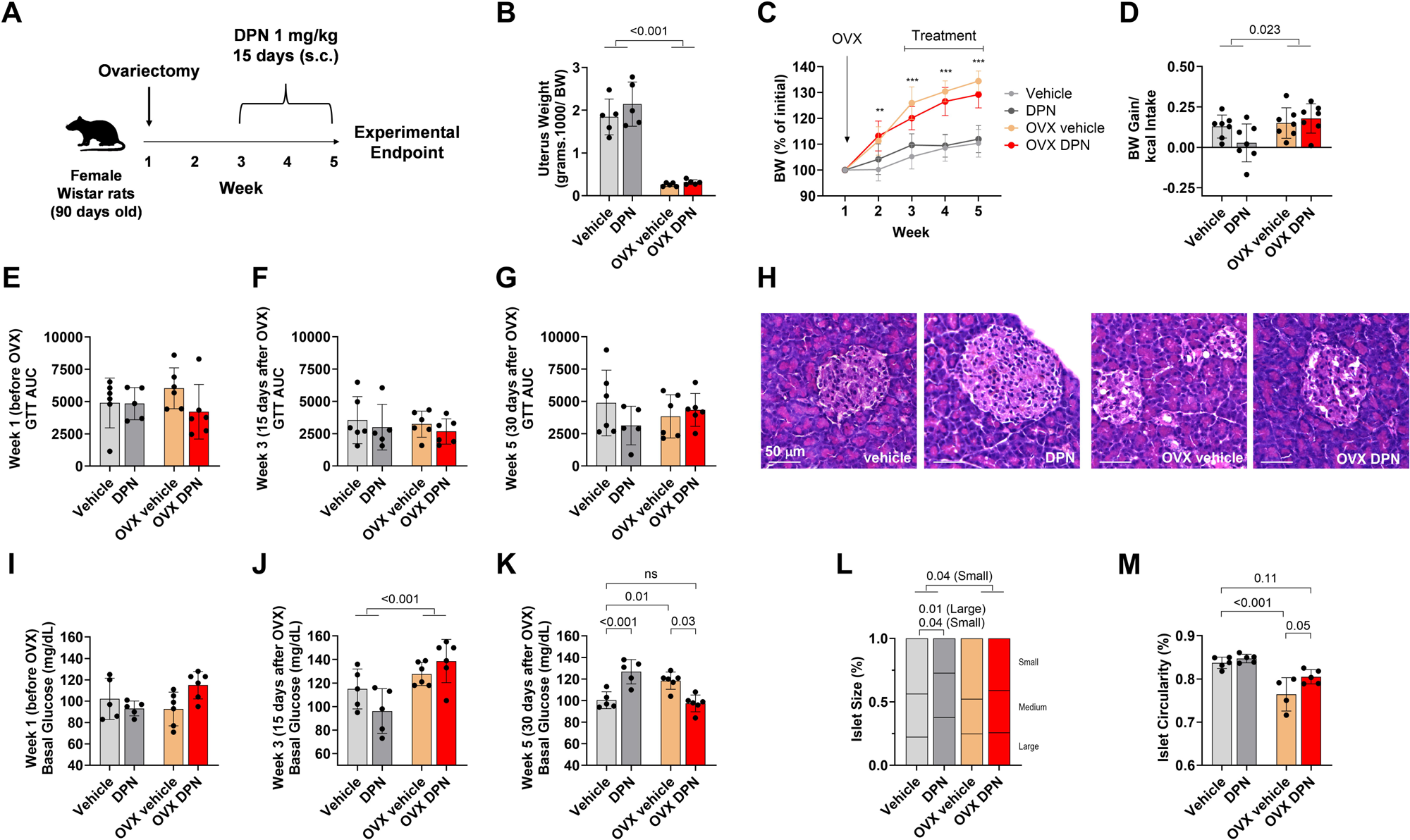
ERβ agonist modulates glucose homeostasis in ovariectomized (OVX) female Wistar rats. Experimental protocol (**A**). Uterus weight (**B**), body weight gain (**C**), final body weight gain per caloric intake (**D**). Week 1 (**E**), week 3 (**F**), week 5 (**G**) GTT AUC. Week 1 (**H**), week 3 (**I**), week 5 (**J**) basal glucose levels. Representative islet histology (**K**), pancreatic islet size distribution (**L**), pancreas circularity (**M**). Groups were compared by two-way ANOVA followed by Tukey posttest. N = 5-6 (A-J) n = 5 (K-M). Significances were considered at p < 0.05.

**Figure 1B** shows that OVX decreases endpoint uterus weights, as expected, with no noticeable effect of DPN. Also, in control animals the estrous cycling was not altered during DPN treatment (**Supplementary material Figure S1**). Weight gain over time (**Figure 1C**) was observed in all groups, and was significantly higher in OVX groups, as expected, with no significant effect of DPN. OVX slightly increased weight gain per calorie ingested, also with no effect of DPN (**Figure 1D**).

Glucose tolerance test (GTT) profiles before treatments at Week 1 (**Figure 1E**, area under the curve, and **Figure 1I**, basal glucose) were similar for all groups, indicating appropriate individual distribution prior to interventions. At Week 3 and 5, GTT response remained unchanged (**Figure 1F and 1G**), although basal glucose levels were increased 15 days after OVX (**Figure 1J**) and remained significantly elevated in OVX animals relative to controls at Week 5 (**Figure 1K**). DPN promoted an increase in basal glucose levels in control animals, but significantly decreased the elevated basal glucose levels observed in OVX rats, reverting them to control levels. This indicates the ERβ agonist promotes significant glycemic modulation in OVX animals.

Given these effects on glucose homeostasis, we also examined endocrine pancreas histology (**Figure 1H**). OVX promoted an increase in smaller islets (**Figure 1L**) and a decrease in islet circularity (**Figure 1M**). DPN specifically increased islet size in the absence of OVX (**Figure 1L**), while partially protecting against the decrease in circularity promoted by OVX (**Figure 1M**).

### 3.2. ERβ activation improves lipid homeostasis in OVX animals

Retroperitoneal white adipose tissue (WATr) was also evaluated in response to OVX and DPN (**Figure 2A**), given its importance as a metabolic hub. As expected, OVX animals had a significant increase in WATr tissue weight, an effect that was abrogated by DPN treatment (**Figure 2B**). Interestingly, DPN also decreased average adipocyte areas in both control and OVX animals (**Figure 2C**).

**Figure 2.**
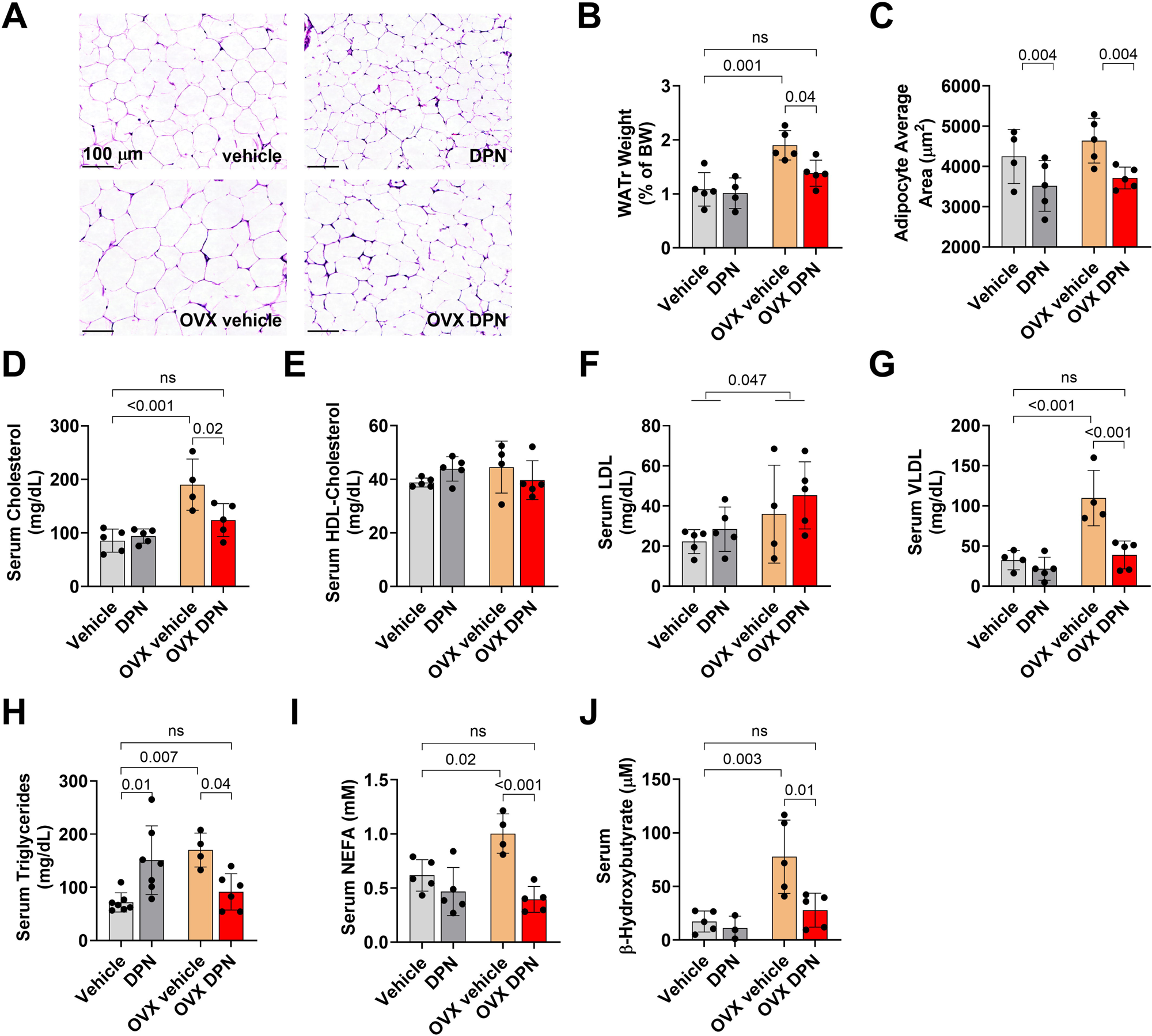
ERβ agonist modulates lipid homeostasis in ovariectomized (OVX) female Wistar rats. Representative WATr histology (**A**). Retroperitoneal white adipose tissue (WATr) weight (**B**), WATr adipocyte area (**C**), serum cholesterol (**D**), HDL (**E**), LDL (**F**), VLDL (**G**), triglycerides (**H**), non-esterified fatty acids (**I**), β-hydroxybutyrate (**J**). Groups were compared by two-way ANOVA followed by Tukey posttest. N = 5. Significances were considered at p < 0.05.

In line with these anatomical findings, the serum lipid profile changed remarkably after OVX. Total cholesterol (**Figure 2D**), LDL (**Figure 2F**), and VLDL (**Figure 2G**), but not HDL (**Figure 2E**), were increased by OVX, with strong protective effects of DPN on total (**Figure 2D**) and VLDL (**Figure 2G**) cholesterol levels. Serum triglycerides were also increased in OVX animals, in a manner prevented by DPN (**Figure 2H**). Serum non-esterified free fatty acid (NEFA) concentrations increased in OVX (**Figure 2I**), an effect accompanied by increased ketone bodies (**Figure 2J**). DPN treatment restored these parameters in OVX animals to control levels (**Figures 2I** and **2J**). Overall, these results indicate that ERβ activation has protective effects against OVX-induced dyslipidemia and also promotes changes in lipid metabolism in the liver, from where ketone bodies are derived.

### 3.3. Post-OVX liver metabolism is modulated by selective ERβ activation

Due to its central function coordinating carbohydrate and lipid metabolism, along with the observed evidence of changes in liver-derived ketone bodies, hepatic tissue was evaluated in more detail. Histological (**Figure 3A**) and content analysis showed increased glycogen in OVX animals, an effect reversed by DPN (**Figure 3B**). Surprisingly, DPN had the opposite effect in non-OVX rats, increasing glycogen content in control animals (**Figure 3B**). Liver triglyceride content was also increased by OVX, but DPN treatment did not alter this content significantly (**Figure 3C**). Consistently, macrovesicular steatosis was augmented by OVX, also with no effect of ERβ activation promoted by DPN (**Figure 3D**). These effects were not accompanied by significant effects in isolated liver mitochondrial Complex I and Complex II-linked oxygen consumption rates, nor changes in H_2_O_2_ production (**Supplementary material Figures S3 and S4**).

**Figure 3.**
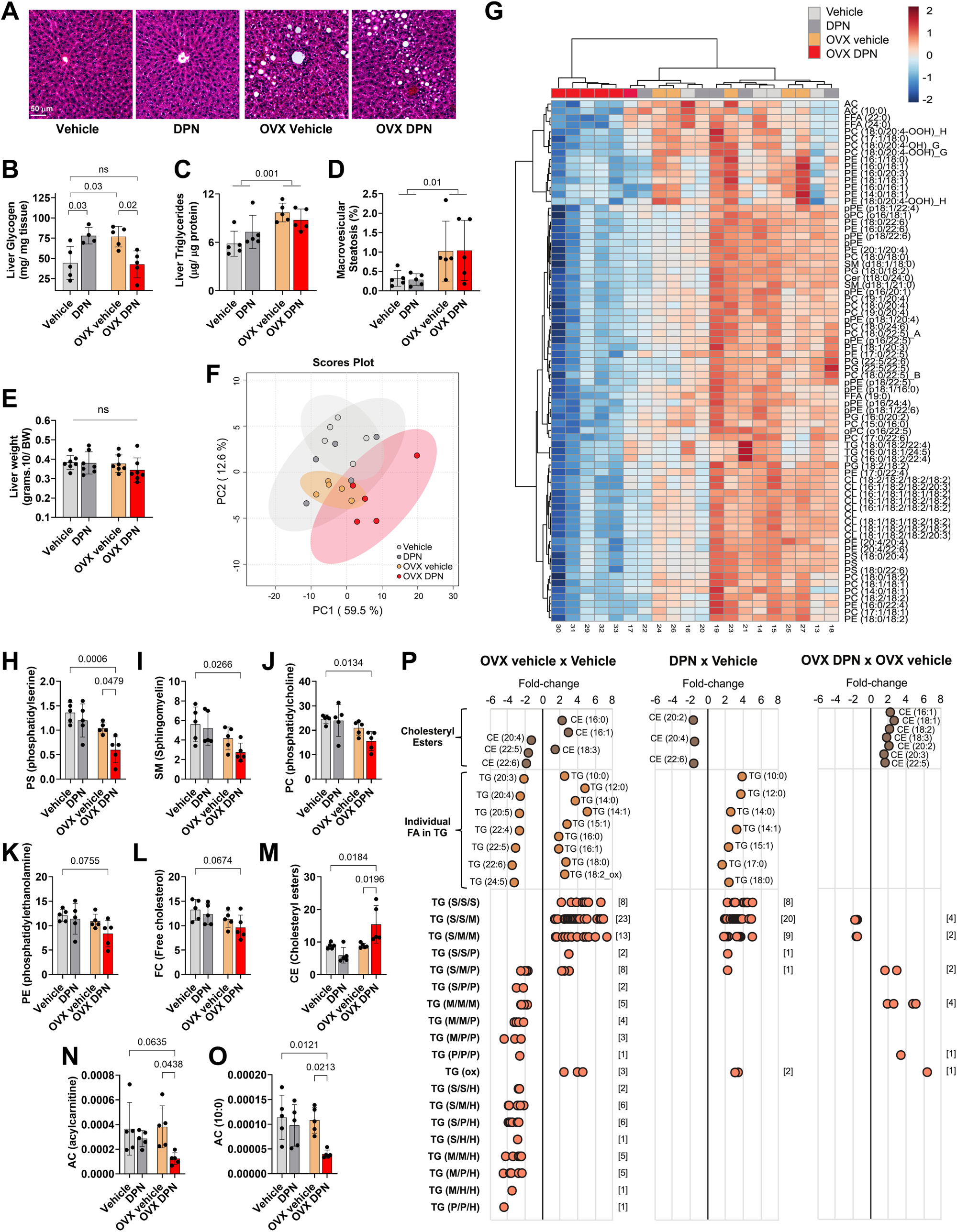
Post-OVX liver metabolism is modulated by selective ERβ activation. Representative liver histology (**A**), liver glycogen (**B**), triglycerides (**C**), macrovesicular steatosis (**D**), liver weight (**E**). Lipidomic Principal Component Analysis (**F**) clustering (**G**), and quantification of phospholipids (**H-K**), free cholesterol (**L**), cholesteryl esters (**M**), acylcarnitine (**N** and **O**), CE and TG associated lipids (**P**) were assessed. Groups were compared by two-way ANOVA followed by Tukey posttest. N = 5 (A-D) n = 6-7 (E-J). Significances were considered at p < 0.05

While total lipid content remained unaffected by the treatment, lipidomic profiling showed that DPN treatment reshaped the lipid composition of OVX animals, resulting in a clearer separation between OVX DPN and OVX vehicle groups in the clustering and PCA analyses, meaning that DPN treatment induced a clear separate clustering of the OVX group (**Figure 3F and G**). DPN promoted a decrease in phospholipids (**Figure 3H-K**) and free cholesterol (FC) (**Figure 3L**) and an increase in cholesteryl esters (CE) (**Figure 3M**). Total acylcarnitines (AC) were also decreased, including AC (10:0), a medium-chain acylcarnitine (**Figure 3N and O**). Analyzing CE species and individual fatty acid (FA) chains in triglycerides uncovered that OVX promoted an increase in saturated and monounsaturated CE and TG fatty acids, with a decrease in polyunsaturated and highly unsaturated FAs (**Figure 3P**). In OVX animals, DPN increased mono, poly and highly unsaturated CE species, while exerting a modest effect on TG species. Overall, these findings indicate that DPN promotes significant lipidome remodeling specific to OVX animals.

### 3.4. Selective ERβ activation remodels hepatocyte lipid profiles *in situ*

Overall, results in the animals suggest that OVX promotes large metabolic modifications, resulting in marked unfavorable changes in circulating lipids, liver lipids, and hepatic-derived ketones, in a manner prevented by DPN. This effect was not explained by a direct effect on mitochondria studied in isolation, as assessed in organelles from the tissue of the treated animals (**Supplementary material Figures S3 and S4**). However, these outcomes could be related to alterations in mitochondrial metabolism *in situ* which depend on cytosolic signaling, and that would not be conserved in experiments with isolated mitochondria. To overcome this limitation, we established a cell model using hepatocytes (AML-12 cells) in which key metabolic features associated with OVX were mimicked by submitting the cells to high glucose combined with palmitate, and the effects of selective ERβ activation were verified by treatment with 0.1 or 1 nM DPN (**Figure 4**).

**Figure 4.**
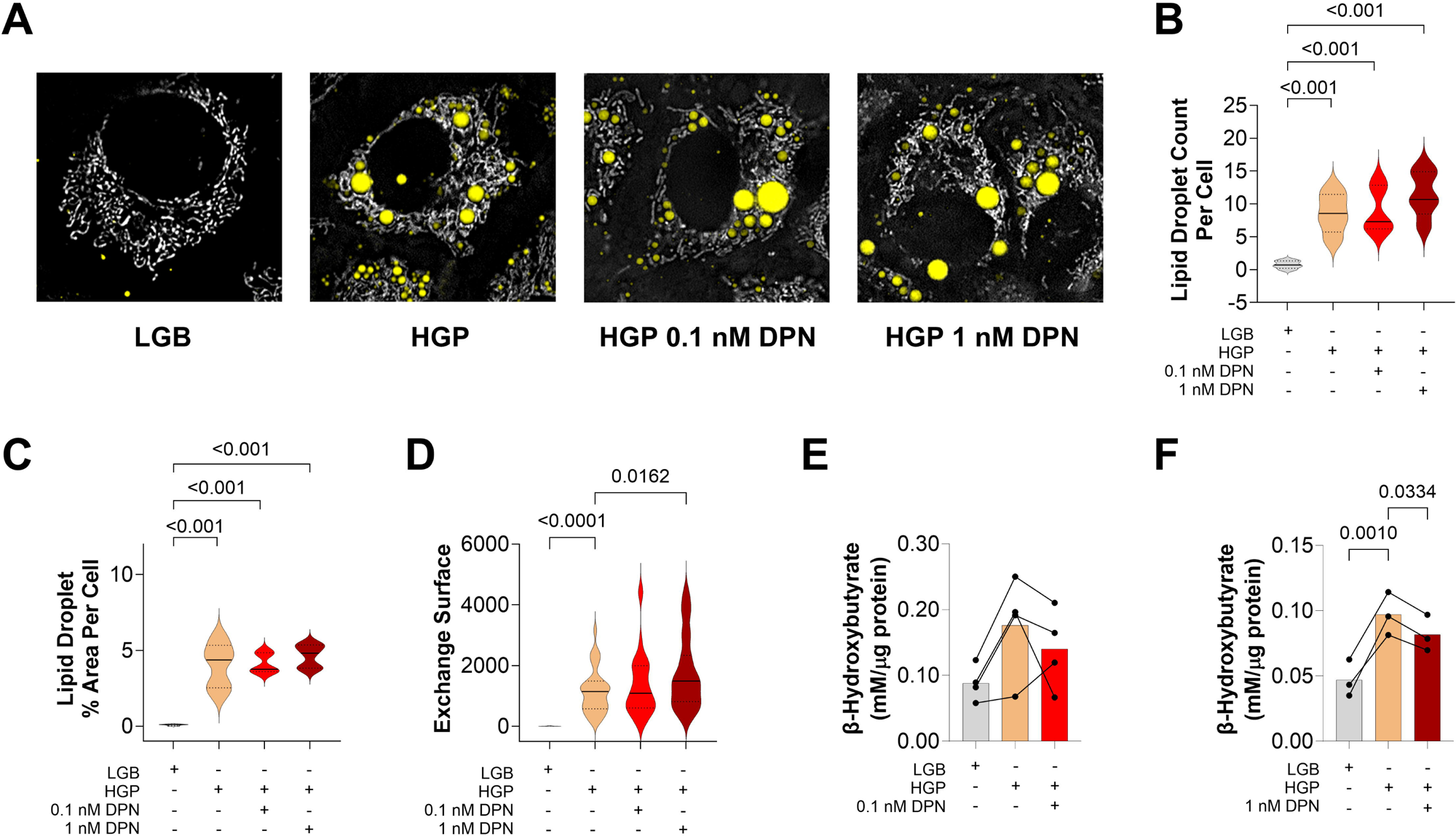
Selective ERβ activation remodels hepatocyte lipid profiles *in situ*. Representative images of mitochondrial network and lipid droplets using Mitotracker deep red (gray) and BODIPY (yellow) (**A**); lipid droplets count (**B**) and area (**C**) per cell and Exchange surface (**D**). β-hydroxybutyrate release to the culture medium after 24 h of treatment (**D** and **E**). Groups were compared by one-way ANOVA. n = 4-5. Significances were considered at p < 0.05.

The exposure of hepatocyte cells to high glucose and palmitate (HGP) led to the accumulation of lipid droplets (LD; **Figure 4A**), with higher numbers of LDs (**Figure 4B**) and higher area of LDs per cell (**Figure 4C**), in a manner unaffected by the presence of DPN. This result mirrors effects of DPN on OVX rat livers. Using maximal projection, we quantified LD surface area (exchange surface, **Figure 4D**), and found it significantly increased after DPN treatment, suggesting an effect of DPN on the LD profile. Furthermore, HGP cells presented increased ketone secretion, in a manner prevented by DPN (**Figure 4E** and **4F**), once again mirroring effects seen in OVX animals. Thus, the HGP hepatocyte model presents similar effects to OVX and responds equally to ERβ activation, validating its use in uncovering mechanisms involved in this metabolic modulation, and indicating that the drug acts directly on liver cells.

We then conducted extracellular metabolic flux analysis in these hepatocytes using the Seahorse system, quantifying basal (**Figure 5A**), ATP-linked (Oligomycin-inhibited, **Figure 5B**), and maximal (Uncoupler-induced, **Figure 5C**) oxygen consumption rates under typical “Mitochondrial Stress-Test” conditions in which glucose is used as the main added extracellular substrate. All these parameters were decreased in cells submitted to HGP relative to low glucose controls (LGB), with a tendency toward some reversal of these changes by DPN, although results were not consistent in all concentrations and conditions.

**Figure 5.**
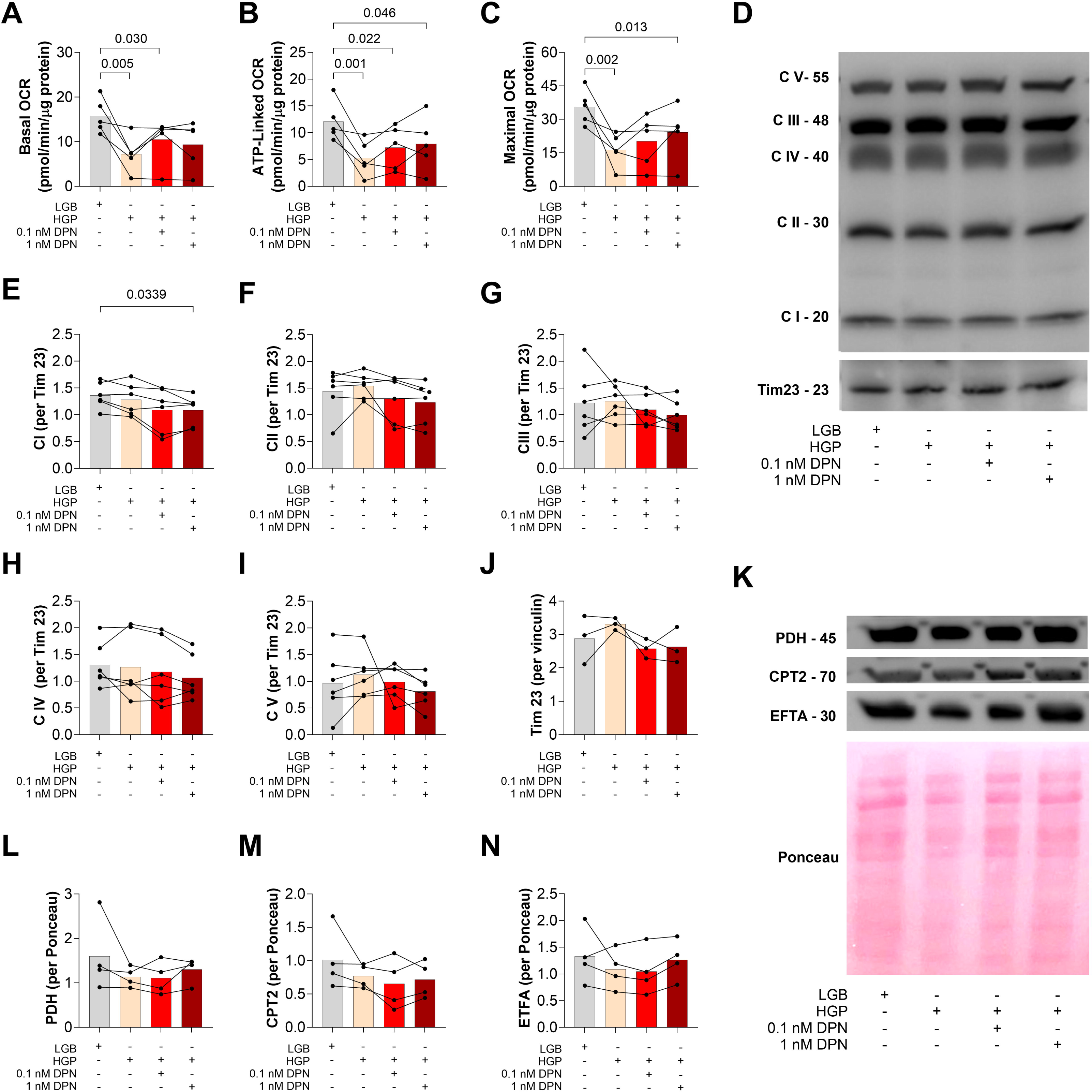
ERβ agonist effects on oxygen consumption, levels of mitochondrial oxidative phosphorylation components and metabolic checkpoints. Basal (**A**), ATP-linked (**B**), and maximal (**C**) oxygen consumption rates. Representative blot of oxidative phosphorylation components using total Oxphos antibody mix (**D**) and TIM23 as internal control. Quantification of protein levels of mitochondrial complexes I, II, III, IV, and V (**E-I**). Representative blots (**K**) and quantifications of pyruvate dehydrogenase (PDH, **L**), Carnitine palmitoyltransferase II (CPT2, **M**), electron transfer flavoprotein subunit alpha (EFTA, **N**). Ponceau staining was used for normalization in L-M. Groups were compared by one-way ANOVA. n = 3-5. Significances were considered at p < 0.05.

We then quantified components of the oxidative phosphorylation machinery through western blot analysis using the Total OxPhos antibody mix (a typical blot is shown in **Figure 5D**). With the exception of CI, OXPHOS complexes as well as inner mitochondrial membrane marker protein Tim 23 did not change in HGP, nor with DPN treatment (**Figure 5E-J**). Interestingly, DPN decreased Complex I protein levels (**Figure 5E**). Pyruvate dehydrogenase (PDH, **Figure 5K** and **5L**) levels remained unchanged, a result that does not explain the increase in oxygen consumption rates promoted by this drug. When looking at other metabolic checkpoints, we found that and DPN did not significantly alter carnitine palmitoyltransferase 2 (CPT2, **Figure 5M**) or electron transfer flavoprotein subunit alpha (ETFA, **Figure 5N**) levels, showing that does not affect quantities of components of fatty acid oxidation. These results indicate that the changes in metabolic fluxes in the presence of DPN involve modulation of metabolic rates rather than control of the amounts of enzymes at the metabolic checkpoints involved.

### 3.5. ERβ selective activation modulates substrate dependence and metabolic flexibility in hepatocytes

We saw changes in lipid composition promoted by DPN both *in vitro* and *in vivo*, but no overt changes in mitochondrial oxygen consumption nor composition compatible with changes in metabolism promoted by DPN. Indeed, while the “Stress Test” conducted in Figure 5 can give important information regarding metabolic state, it does not typically include added fatty acids as substrates, nor measure specific substrate oxidation capacity, dependency or flexibility.

Extracellular flux measurements can be designed to evaluate the participation of different metabolic pathways in intact cells by following the effects of inhibitors of different metabolic pathways, alone or in combination. We used UK5099, a mitochondrial pyruvate carrier inhibitor, to specifically inhibit complete glucose oxidation (generating pyruvate, which enters mitochondria), Etomoxir (a carnitine palmitoyl transferase inhibitor) to prevent uptake of fatty acids into mitochondria, and BPTES, a glutaminase inhibitor, to assess the roles of carbohydrate, fatty acid, and amino acid oxidation on metabolic fluxes in hepatocytes (**Figure 6A**). With sequential additions of the inhibitors alone or in combination, the dependence and capacity of each pathway can be assessed: <u>dependence</u> is evaluated by inhibiting the target pathway first, then the other pathways, while <u>capacity</u> is evaluated by inhibiting alternate pathways first, followed by the target pathway (**Figure 6A**). Finally, metabolic <u>flexibility</u> can be calculated as the difference between dependence and capacity. Experiments were done in parallel for two concentrations of DPN (0.1 and 1 nM) due to the number of samples needed, in order to conduct controls in parallel for the same sample.

**Figure 6.**
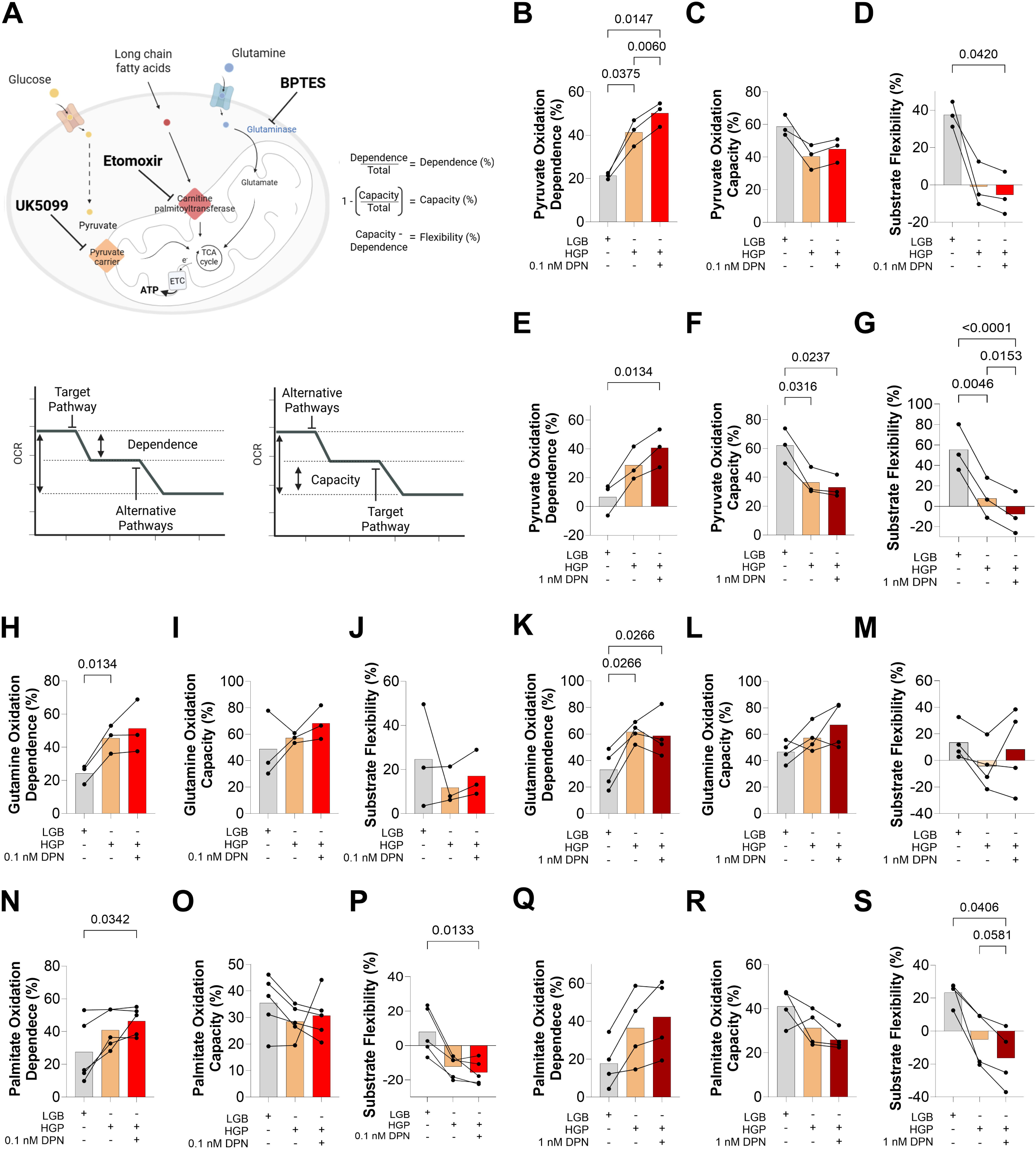
ERβ activation modulates substrate oxidation dependence and flexibility under conditions of energy overload. Oxygen consumption rates (OCR) were measured under basal conditions, and after the injection of 7 μM Etomoxir (CPT1 inhibitor), 6 μM UK5099 (mitochondrial pyruvate carrier inhibitor) and/or 8 μM BPTES (glutaminase inhibitor), as indicated, to measure dependence and capacity (**Panel A**, adapted from Agilent (76), created with Biorender). Capacity was measured after injecting all inhibitors except for the target oxidative pathway inhibitor. Dependence was expressed as the target substrate per all inhibitor response ratio. Capacity was expressed as the alternative substrates’ response per all inhibitors’ response ratio. Flexibility is the difference between capacity and dependence. Values were quantified for the different substrates (Pyruvate, **B-G**; Glutamine, **H-M**; and Palmitate, **N-S**) and using two DPN concentrations, as shown. Groups were compared by one-way ANOVA. n = 3-4. Significances were considered at p < 0.05.

Overall (**Figure 6B-S**), we found that nutrient overload with the HGP protocol did not increase the capacity to oxidize pyruvate, glutamine, or palmitate relative to low glucose (LGB) conditions (panels **6C**, **6F**, **6I**, **6L**, **6O**, and **6R**). Conversely, the dependence on the oxidation of glutamine (**Figures 6H** and **6K**) and pyruvate (**Figures 6B** and **6E**) increased under HGP conditions, leading to a reduced metabolic flexibility (**Figures 6D**, **6G**, **6J** and **6M**). Interestingly, by significantly increasing dependence (**Figures 6B**, **6E**, **6N** and **6Q**) on palmitate (**Figures 6P** and **6S**) and pyruvate (**Figures 6D** and **6G**), DPN further reduced flexibility on these substrates.

Together, these metabolic flux quantifications demonstrate that selective ERβ activation by DPN in hepatocytes promotes complete mitochondrial oxidation of carbohydrate-derived pyruvate and fat-derived palmitate, a result compatible with the fact that it decreases VLDL, triglycerides, fatty acids, and ketones *in vivo*, and promotes lipid droplet remodeling with enhanced exchange surface *in vitro*.

## 4. DISCUSSION

Estrogens modulate growth and function of the reproductive tissues, effects which can also be achieved in postmenopause by treatment with classical estrogen receptor agonists. Non-selective estrogen receptor agonists also have metabolic effects, at least partially reversing the undesirable metabolic impacts of menopause (8–10). However, these beneficial results are often unachievable in post-menopausal women who have predisposition toward hormone-responsive cancers, and therefore are not candidates for classic hormone replacement therapy. In this sense, it is of interest to understand the metabolic impact of selective ERβ activation, which is not associated with increased cancer risk (23–28). Fittingly, we found that the *in vivo* treatment with the selective ERβ agonist DPN had no change in wet uterus weights, a result in line with existing literature suggesting that ERβ does not exert trophic effects on the reproductive organ in rodents (44,45), nor influence the hypothalamic-pituitary-gonadal axis (46).

A well-documented consequence of estradiol deficiency on energy metabolism is persistent weight gain, not always correlated with food intake (20,47,48). Indeed, we find that OVX animals exhibited marked weight gain beginning the first week post-ovariectomy, which remained pronounced until the experimental endpoint (30 days). Body weight gain per caloric intake was elevated in OVX animals, indicating increased energy conversion efficiency induced by estradiol deficiency. Selective ERβ activation with DPN had no effect on body weight gain or weight gain relative to caloric intake in OVX animals, a finding consistent with a prior study showing that ERβ agonists do not significantly affect body weight (17).

Although the ERβ agonist did not influence body weight, marked changes were observed in glucose and lipid metabolism and content after treatment was introduced post-OVX. OVX induced enhanced basal glucose levels, accompanied by changes in pancreatic islet size and circularity. OVX also increased WATr weight and elevated serum cholesterol, LDL, VLDL, triglycerides, and fatty acids - confirming the establishment of an energy overload state induced by estradiol deficiency, characterized in previous literature by elevated fasting glucose, development of insulin resistance (20), increased white adipose tissue mass, and a disrupted circulating lipid profile (48). DPN treatment did not alter glucose tolerance assessed in GTT, but restored basal glucose levels in OVX animals to those observed in controls. DPN also improved islet circularity, preserving physiological pancreatic islet morphology. Moreover, treatment with DPN decreased WATr weight and adipocyte areas in both control and OVX animals, accompanied by improvements in serum cholesterol, VLDL, triglycerides, and FFAs. These data demonstrate that selective ERβ activation, even after estradiol deficiency establishment, can restore critical markers of metabolic disease risk, improving both glucose and lipid homeostasis, in many cases to levels indistinguishable from non-OVX controls.

The effects on glucose homeostasis and islet morphology suggest DPN may also exert direct actions on the endocrine pancreas. *In vitro* studies suggest that ERβ, along with ERα and GPER, enhances β-cell survival and function, with ERβ contributing toward apoptosis prevention and K_ATP_ inhibition mimicry (49,50). ERβ^−/−^ mice showed delayed first-phase insulin release following glucose stimulation, but with no changes in basal glucose levels (51). Another study found that DPN did not significantly affect glucose or glycated hemoglobin (HbA1c) in OVX Sprague-Dawley rats (52). Conversely, an ERβ agonist decreased serum glucose levels and improved β-cell function in high fat-fed mice (53), leading to the conclusion that the direct effects of the receptor on β-cells seem to be dependent on the metabolic state and animal model.

Visceral adiposity reduction may explain decreased circulating FFAs promoted by DPN treatment observed in our *in vivo* model, as visceral fat exhibits high lipolytic activity, linked to a detrimental cardiovascular profile and β-cell dysfunction (54–56). Indeed, ERβ activation may promote adipose tissue remodeling (17). In male rats, ERβ agonist treatment shifted fat distribution from visceral to subcutaneous without affecting body weight - a pattern associated with reduced lipolytic activity and improved insulin sensitivity (17). In OVX females, ERβ agonists prevented WAT gain, enhanced lean mass (57), reduced LDL, and improved hepatic insulin sensitivity under high-fat diet conditions (53). We found that serum β-hydroxybutyrate levels followed the same pattern as FFAs - increased in OVX animals and fully restored to control levels after DPN treatment. As incomplete lipid oxidation to acetyl CoA is associated with hepatic ketone body production (58,59), these findings implicate the liver, alongside the adipose tissue and endocrine pancreas, in the beneficial metabolic effects observed with ERβ activation *in vivo.* Given the liver’s central role in metabolism, and the scarcity of ERβ-focused liver studies, we investigated hepatic metabolic status in OVX females.

We found that OVX induced significant changes in liver metabolism, with increased glycogen and lipid accumulation, suggesting substrate overload and enhanced storage signaling. Notably, the GTT curve did not differ between OVX and control animals, suggesting a similar insulin sensitivity. However, increased ketone production alongside lipid accumulation suggests early-stage metabolic-associated fatty liver disease (MASLD)-like decompensation (60,61), although macrovesicular steatosis in our samples remained within physiological levels (39). In line with this, the increase in monosaturated and the decrease in highly unsaturated cholesteryl ester accompanied by the same pattern in triglyceride fatty acid species in OVX (62) indicates an ongoing dysregulation of hepatic lipid processing. Interestingly, DPN treatment promoted a lipidome shift toward a protective profile, characterized by decreased FFAs and increased esterified cholesterol ester species, particularly those containing monounsaturated, polyunsaturated, and highly unsaturated fatty acids. This aligns with literature showing ERβ agonists reduce saturated fatty acids while increasing unsaturated species in high-fat diet models of non-ovariectomized females and in obese male mice (17,53), and shows the potential of ERβ as a target in metabolic overload under estradiol-deficient conditions.

We also assessed mitochondrial function in freshly isolated liver mitochondria. Both state 3 (phosphorylating) and state 4 (proton-leak dependent) oxygen consumption were elevated in St3 consumption (CI and CII-linked) in OVX animals, indicating higher electron transport rates and, but reduced coupling efficiency (CII-linked) between oxygen consumption and ATP synthesis (**Figure S3**). This inefficiency is a known feature of estradiol deficiency and is recoverable via non-selective estradiol treatment (63,64), implicating estrogen receptors in mitochondrial regulation (64–66). Fittingly, our results showed that DPN-treated OVX animals did not have differences in mitochondrial oxygen coupling rate when compared to controls, indicating that the treatment restores these oxidative phosphorylation parameters. These results are different from previously reported estradiol (non-selective) effects (64–66) and suggest ERβ may have distinct impacts on mitochondrial respiration.

While ERβ mitochondrial effects in liver had not been uncovered previously, some work has shown that ERβ appears to be involved in hepatic lipid metabolism, and may attenuate MASLD progression (57,67), despite its lower expression levels (22,57) in this organ. Our findings indicate that OVX induced lipid accumulation and mild steatosis without structural liver damage, indicating subclinical metabolic imbalance. DPN treatment did not alter total liver lipid quantities nor histological macrovesicular steatosis, but profoundly shifted lipid species composition. ERβ activation thus triggers a lipidomic shift that favors a protective profile, with the conversion of toxic free fatty acids into triglycerides, acting as a buffering mechanism that correlates with improved serological outcomes (68,69).

To understand the changes in metabolism happening in liver cells in terms of specific metabolic modulation, we also conducted *in vitro* studies with hepatocytes. Our model reproduced the effects of nutrient overload seen *in vivo*, with accumulation of lipid droplets and enhanced ketone release. As seen *in vivo*, DPN prevented ketone production (70,71), but not lipid accumulation. A notable finding *in vivo* was the decrease in several phospholipid species, which can be associated with the presence of supersized lipid droplets (LDs), that have lower exchange surfaces and may be less accessible for metabolization, thereby impairing lipid oxidation (72,73). Interestingly, quantification *in vitro* revealed that the DPN treatment increased exchange surface area of the LDs, suggesting enhanced accessibility for metabolization. Overall, our data show that ERβ activation directly regulates the specific types of FFAs available for oxidation within hepatocytes. By modulating this lipid supply, the treatment prevents acetyl-CoA overload, the primary substrate for ketogenesis. As effects can be reproduced in liver cells, these metabolic shifts are driven by direct cellular mechanisms rather than by systemic effects of the drug.

Interestingly, estradiol is known to enhance hepatic oxidative machinery and substrate oxidation (74,75), a result that is partially compatible with our results suggesting ERβ may promote increased dependence and decreased flexibility of full oxidation of carbohydrate-derived pyruvate and the fatty acid palmitate. This switch promotes complete acetyl-CoA oxidation, accompanied by decreased long-chain acylcarnitines, explaining reduced incomplete lipid oxidation and ketone body release under nutrient overload seen in both our *in vitro* and *in vivo* models (74).

Overall, we find that ERβ activation promotes specific metabolic reconfiguration in liver cells that impact on *in vivo* lipid and glucose metabolism, promoting improved systemic glucose and lipid homeostasis as well as positively modulating the liver lipidome, despite persistent energy overload seen in the absence of estrogens. These findings highlight ERβ activation as a promising alternative to non-selective estradiol therapy, particularly for postmenopausal women with established metabolic imbalances.

## ABBREVIATIONS

AC: acylcarnitine
AML-12: alpha mouse liver 12 cells
BSA: bovine serum albumin
BW: body weight
CCCP: carbonyl cyanide m-chlorophenyl hydrazone
CE: cholesteryl esters
CPT2: carnitine palmitoyltransferase 2
DMEM: Dulbecco's modified Eagle's minimal essential medium
DPN: diarylpropionitrile
ER: estrogen receptor
ER?: estrogen receptor alpha
ER?: estrogen receptor beta
ER: estrogen receptor beta
ETFA: electron transfer flavoprotein subunit alpha
FA: fatty acid
FBS: fetal bovine serum
FC: free cholesterol
FFA: free fatty acid
GPER: G protein-coupled estrogen receptor
GTT: glucose tolerance test
HDL: high-density lipoprotein
HGP: high glucose-palmitate
HRT: hormone replacement therapy
LD: lipid droplet
LDL: low-density lipoprotein
LGB: low glucose-BSA
MTBE: methyl-tert-butyl ether
NEFA: non-esterified free fatty acid
OCR: oxygen consumption rate
OVX: ovariectomy
PCA: principal component analysis
PDH: pyruvate dehydrogenase
PKA: protein kinase A
PKC: protein kinase C
TG: triglycerides
Tim23: translocase of the inner mitochondrial membrane 23
VLDL: very low-density lipoprotein
WAT: white adipose tissue
WATr: retroperitoneal white adipose tissue.

## ACKNOWLEDGEMENTS

The authors acknowledge Professor Vilma Regina Martins for intellectual contributions. We also thank Silvânia M. P. Neves and animal facility crew for excellent animal care, Sirley Mendes de Oliveira for technical support and Mario Costa Cruz (CONFOCAL – CEFAP facility) for histological image capturing.

## DATA AVAILABILITY

Full data has been deposited in Zenodo: 10.5281/zenodo.19487483

## AUTHOR CONTRIBUTIONS

DSR conceived this research, analyzed data, performed experiments, interpreted results, prepared figures, and drafted the manuscript, under supervision of AJK; EAVB, ELV, MICQ, CCCS, MY, and SM performed *in vivo* experiments and interpreted results; MPTC performed *in vitro* experiments and interpreted results. All authors revised and approved the final version of the manuscript.

