## Supplementary material for "Estrogen Receptor Beta Activation Coordinates Liver Lipid Remodeling and Metabolic Fluxes, Preventing Lipotoxicity"

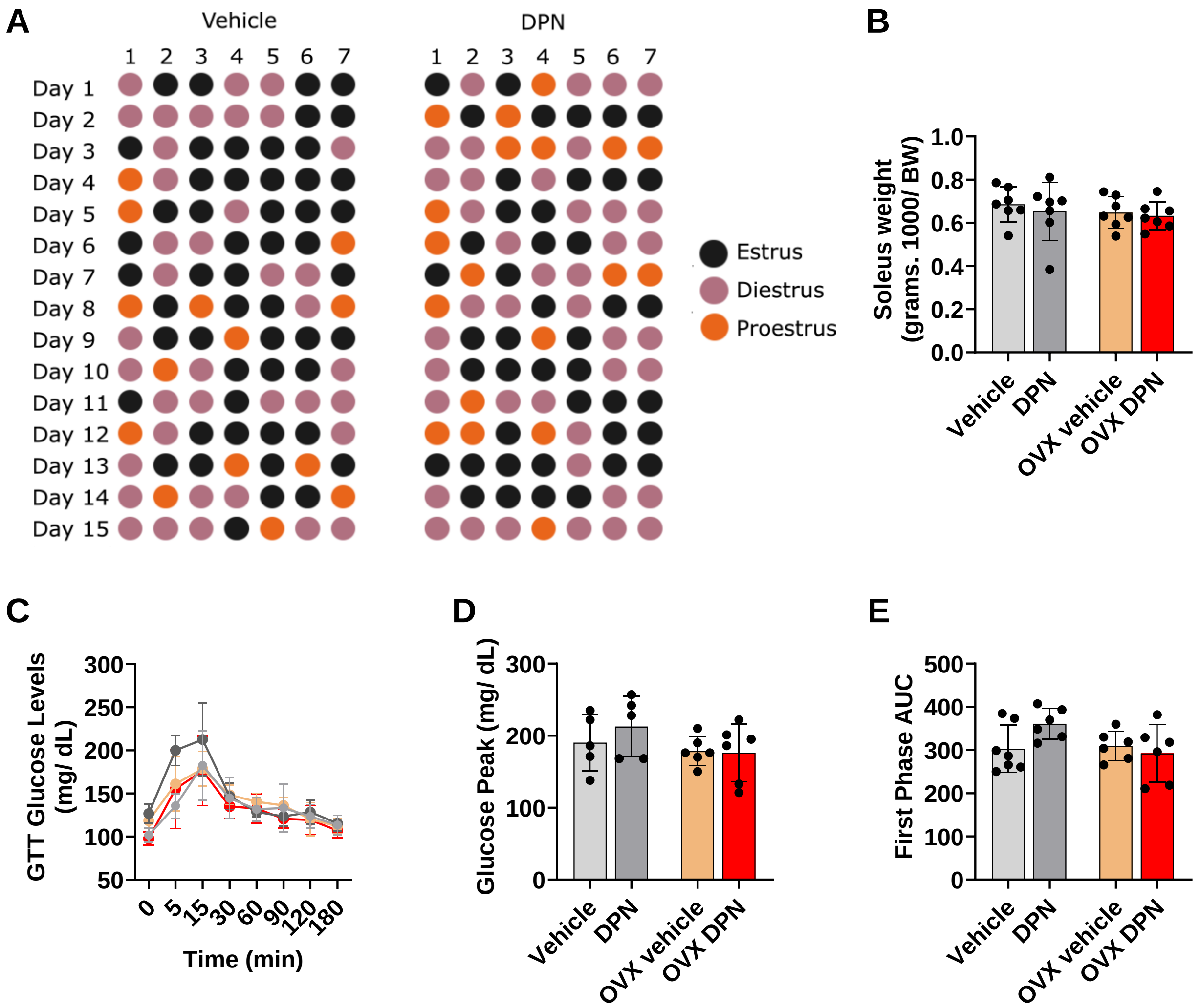


**Figure S1. DPN treatment does not alter estrous cyclicity, muscle weight, or glucose tolerance in intact and OVX animals. (A)** Vaginal smear categorization of 7 individual animals from the Vehicle and DPN groups. **(B)** Soleus weight. **(C)** GTT curve. **(D)** Glucose peak**. (E)** First-phase AUC.

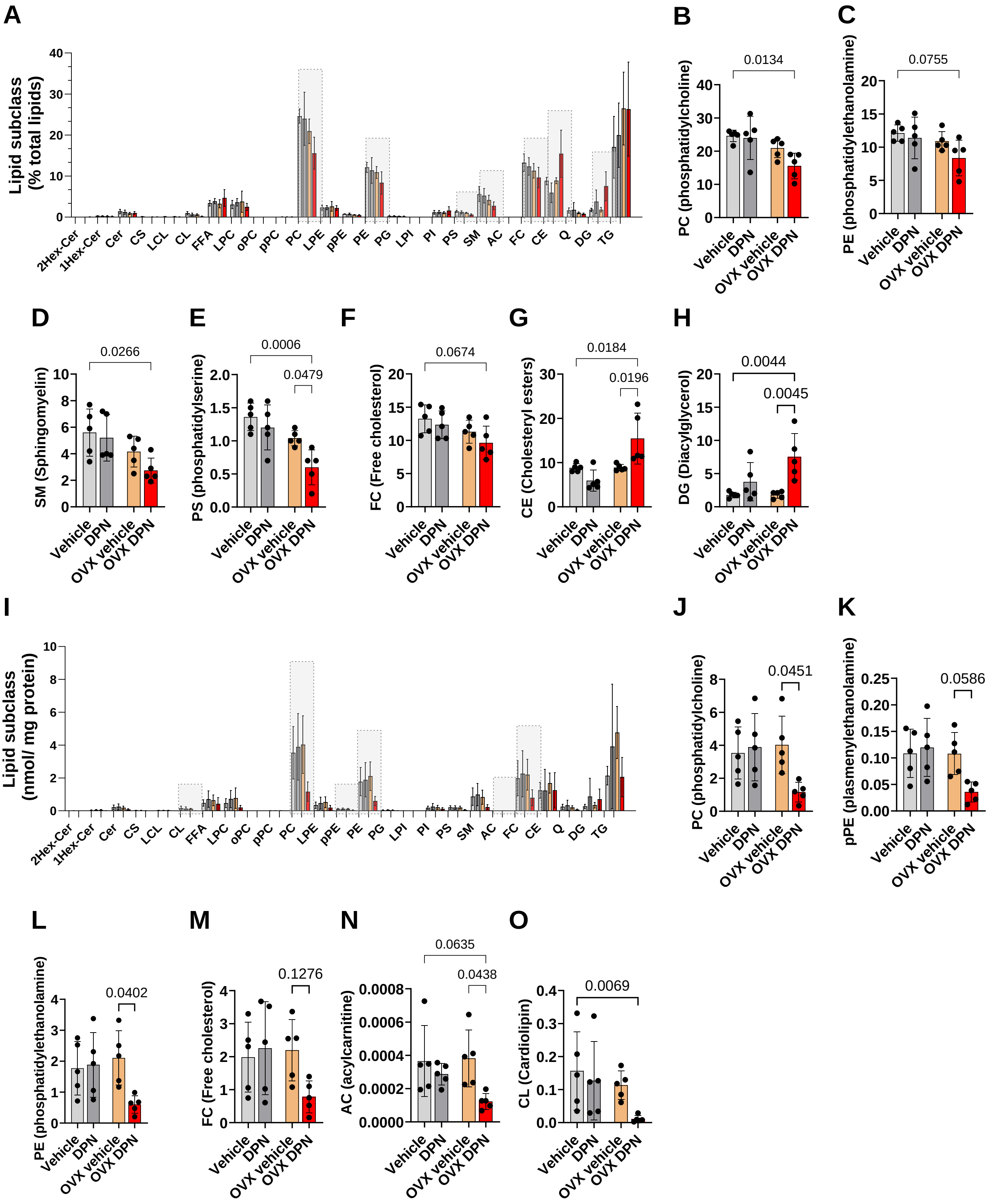
A
Lipid subclass (% total lipids)
2Hex-Cer 1Hex-Cer Cer CS LCL CL FFA LPC oPC pPC PC LPE pPE PE PG LPI PI PS SM AC FC CE Q DG TG
B
PC (phosphatidylcholine)
0.0134
C
PE (phosphatidylethanolamine)
0.0755
D
SM (Sphingomyelin)
0.0266
E
PS (phosphatidylserine)
0.0006
0.0479
F
FC (Free cholesterol)
0.0674
G
CE (Cholesteryl esters)
0.0184
0.0196
H
DG (Diacylglycerol)
0.0044
0.0045
I
Lipid subclass (nmol/ mg protein)
2Hex-Cer 1Hex-Cer Cer CS LCL CL FFA LPC oPC pPC PC LPE pPE PE PG LPI PI PS SM AC FC CE Q DG TG
J
PC (phosphatidylcholine)
0.0451
K
pPE (plasmenylethanolamine)
0.0586
L
PE (phosphatidylethanolamine)
0.0402
M
FC (Free cholesterol)
0.1276
N
AC (acylcarnitine)
0.0635
0.0438
O
CL (Cardiolipin)
0.0069
Vehicle DPN OVX vehicle OVX DPN

**Figure S2. Profile of hepatic lipid subclasses in OVX animals after DPN treatment. (A)** Lipid subclass distribution per group (light grey columns = vehicle, dark grey = DPN, yellow = OVX vehicle, red = OVX DPN), **(B)** Phosphatidylcholine, **(C)** Phosphatidylethanolamine, **(D)** Sphingomyelin, **(E)** phosphatidylserine, **(F)** Free cholesterol, **(G)** Cholesteryl esters, and **(H)** Diacylglycerol (% total lipids). **(I)** Lipid subclass, **(J)** Phosphatidylcholine, **(K)** Phosphatidylethanolamine, **(L)** Plasmenylethanolamine, **(M)** Free cholesterol, **(N)** Acylcarnitine, and **(O)** Cardiolipin (nmol/ mg protein).

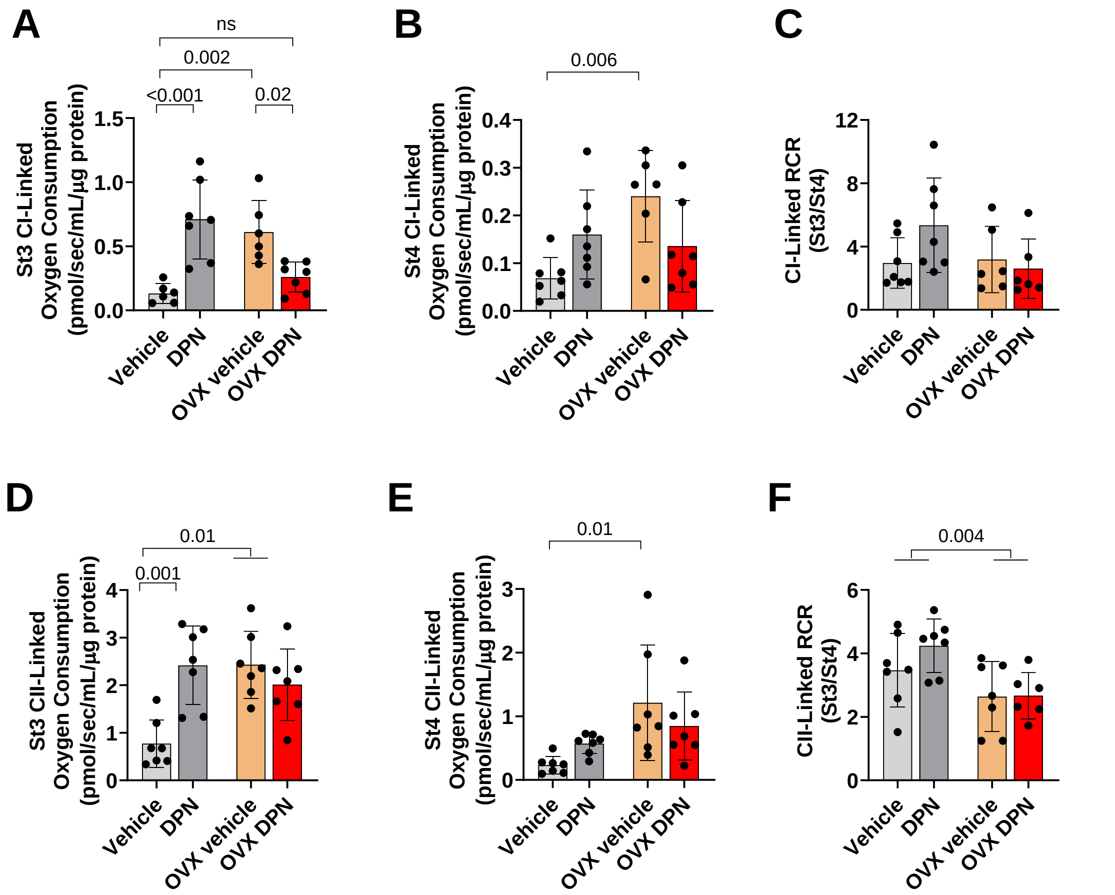


**Figure S3. Oxygen consumption in liver mitochondria from OVX animals after DPN treatment.** **(A)** State 3 Complex I-linked oxygen consumption, **(B)** State 4 Complex I-linked oxygen consumption, **(C)** Complex I-linked Respiratory Control Ratio (RCR)**. (D)** State 3 Complex II-linked oxygen consumption, **(E)** State 4 Complex II-linked oxygen consumption, and **(F)** Complex II-linked RCR.

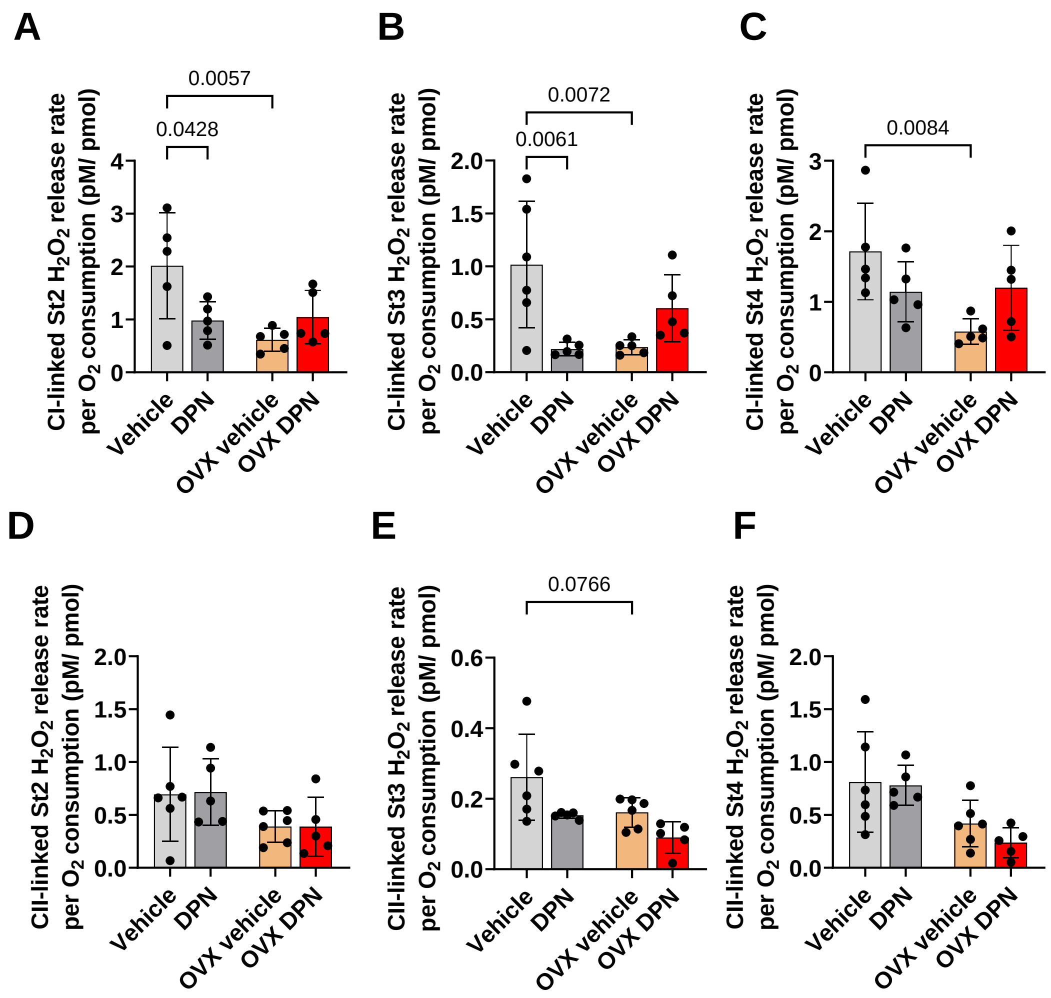


**Figure S4. $H_2O_2$ release rate in liver mitochondria from OVX animals after DPN treatment**. **(A)** basal Complex I-linked $H_2O_2$ release, **(B)** State 3 Complex I-linked $H_2O_2$ release, and **(C)** State 4 Complex I-linked $H_2O_2$ release. **(D)** Basal Complex II-linked $H_2O_2$ release, **(E)** State 3 Complex II-linked $H_2O_2$ release, and **(F)** State 4 Complex II-linked $H_2O_2$ production.
